# Experimental safety testing confirms that the NSAID nimesulide is toxic to *Gyps* vultures in India

**DOI:** 10.1101/2023.06.27.546673

**Authors:** Karikalan Mathesh, Kesavan Manickam, John W. Mallord, K. Mahendran, M. Asok Kumar, Debasish Saikia, S. Chandra Mohan, V Beena, P. Sree Lakshmi, Nikita Prakash, Rohan Shringarpure, Abhijit Pawde, Rhys E. Green, Vibhu Prakash

## Abstract

Population declines of *Gyps* vultures throughout South Asia were caused by unintentional poisoning by the NSAID diclofenac, which was subsequently banned. However, other vulture-toxic NSAIDs are available, including nimesulide, which, in experiments carried out in South Africa, was shown to be toxic to *Gyps* vultures. We report on safety-testing of nimesulide carried out on Himalayan Griffons *G. himalayensis*. We gave two vultures a dose of nimesulide by oral gavage at the maximum level of exposure, with two controls dosed with benzyl alcohol. In the two tested birds, plasma nimesulide concentrations peaked after six hours, while serum uric acid concentrations increased steadily up until 24 hours post-treatment, after which both birds died, displaying severe visceral gout. The control birds showed no adverse clinical or biochemical signs. We confirm that nimesulide is toxic to *Gyps* vultures. Veterinary use of nimesulide should be banned in all *Gyps* vulture range countries in the region.

## 1. Introduction

From being some of the most common large raptors in the world (Houston 1985), three species of *Gyps* vultures endemic to southern Asia were driven to near-extinction, due to unintentional poisoning by the veterinary non-steroidal anti-inflammatory drug (NSAID) diclofenac (Oaks *et al*. 2004, Shultz *et al*. 2004, Green *et al*. 2004, Prakash *et al*. 2007), and all are now classified as Critically Endangered in the IUCN Red List (Birdlife International 2021). In India, the population of the worst-affected species, White-rumped Vulture *G. bengalensis* declined by 99.9% between the early 1990s and 2007, while that of Indian *G. indicus* and Slender-billed Vulture *G. tenuirostris* combined declined by 96.8% (Prakash *et al*. 2007). Vultures ingested diclofenac when they fed on carcasses of domesticated ungulates that had been given the drug shortly before death, a common practice in South Asia. The drug causes kidney failure, elevated plasma uric acid concentrations and death within a few days of ingestion, with severe visceral gout and necrosis of kidney tissues being typical post-mortem signs (Oaks et al. 2004; Swan et al. 2006). Veterinary use of diclofenac was banned in India, Pakistan and Nepal in 2006, and 2010 in Bangladesh (Prakash *et al*. 2012, Sarowar *et al*. 2016), which has contributed to the partial recovery of populations in Nepal (Galligan *et al*. 2020) and the slowing or halting of declines in India and Pakistan (Chaudhry *et al*. 2012, Prakash *et al*. 2017).

However, other NSAIDs are widely available that are also toxic to vultures (Galligan *et al*. 2021). These include ketoprofen, (Naidoo *et al*. 2010), carprofen (Fourie *et al*. 2015), aceclofenac (Galligan *et al*. 2016, Chandramohan *et al*. 2022), and flunixin (Zorilla *et al*. 2015, Herrero-Villar *et al*. 2020). Only two NSAIDs have been proven to be safe for *Gyps* vultures at doses likely to be encountered by wild birds: meloxicam (Swan *et al*. 2006, Swarup *et al*. 2007) and tolfenamic acid (Chandramohan *et al*. 2021). In addition, several dead *G. bengalensis* have been found in India with visceral gout and residues of the NSAID nimesulide (Cuthbert *et al*. 2016; Nambirajan et al. 2021). Subsequently, nimesulide was found to be toxic to *Gyps* vultures (Galligan *et al*. 2022) in experimental tests on captive vultures. In that study, two African Cape Vultures *G. coprotheres* that were given a dose of nimesulide likely to be encountered by vultures in the wild both died from kidney failure. These birds showed similar symptoms to those that had died of diclofenac poisoning (Oaks *et al*. 2004), displaying extensive visceral gout (formation of uric acid crystals, especially in and on the kidney and liver) (Galligan *et al*. 2022). This was accompanied by a 27-79-fold increase in the concentration of uric acid in the blood plasma of the treated birds, which died 27 and 29 hours after treatment.

Here, we report on a study, experimentally testing the toxicity of nimesulide on a near-threatened species of *Gyps* vulture, Himalayan Griffon, *G. himalayensis*, which breeds in mountainous areas of India and neighbouring countries, is a common winter visitor to the north Indian plains and is sensitive to diclofenac nephrotoxicity (Das *et al*. 2010). We followed the protocol of Galligan *et al*. (2022), giving the same dose (per kilogram body weight) of nimesulide, which was based on a vulture’s maximum likely exposure to the drug in the wild, estimated from a pharmacokinetic study of the drug in cattle.

## 2. Methods

### 2.1 Trial animals: housing and management

The experiments were carried out at the Vulture Conservation Breeding Centre at Pinjore, Haryana, India in February 2022. Fifteen wild *G. himalayensis* were trapped on 18^th^ and 25^th^ February in a large (27 × 5 × 9 m) baited walk-in cage trap. They were transferred to a four-compartment purpose-built aviary, each compartment being 6 × 6 × 5 m, and holding a maximum of four vultures. A health check was performed on the birds on 26^th^ February, involving an external examination of body weight, condition and plumage quality, and monitoring of body temperature. Birds were kept in the aviary for three weeks to acclimatize, before the experiment commenced on 11^th^ March. They were fed with 5 kg of goat meat, sourced to be free of NSAIDs, on two days per week, and provided with water *ad libitum*. Samples were taken for haematological and serum analysis (Supplementary Information, Appendix B, Table S1). Surviving birds were released back to the wild on 20^th^ April.

### 2.2 Treatment and study design for oral gavage experiments

Two birds were randomly chosen to be the test subjects and another two acted as controls. The two test subjects were both juveniles, less than one year old; one of the control birds was also a juvenile, the other was in its second year. Before the experiment, we weighed each vulture (treatment group, weight = 8.0 kg for both birds; control group, weight = 8.0 kg and 8.8 kg), and a baseline blood sample taken. We used a commercial injectable brand of veterinary nimesulide (Nimovet, Indian Immunologicals Limited, Hyderabad, India), which is widely available and was purchased from a local pharmacy. The product had a stated nimesulide concentration of 100 mg l^-1^. We administered an aqueous solution of 1% benzyl alcohol and 10% ethanol, which is the carrier solution of nimesulide in the Nimovet formulation, to the control birds by gavage.

We calculated the maximum likely meal weight (*M*) using estimates of the mean daily energy use (*DEU*) of individual vultures. We calculated DEU from the body weight (W = 8.0 kg) of the vulture, *DEU* = 668.4\**W*^0.622^. Meal weight was calculated, as *M* = 3*(*DEU* / 5160), where 5160 kJ kg^-1^ is the energy assimilated by a vulture (Galligan *et al*. 2022), the figure was multiplied by 3 to reflect that the maximum meal size of *Gyps* vultures is typically about three times the amount of food required per day (Swan et al. (2006a). This gave a maximum likely meal of 1.4 kg for the two vultures in the present study. A pharmacokinetics and tissue residue experiment in cattle (Galligan *et al*. 2022) identified the tissue with the highest mean concentration of nimesulide as the injection site muscle at 134.08 mg kg^-1^ (*Rmax*), with the mean weight of that muscle being 1.11 kg (*Vmax*) (Galligan *et al*. 2022). The next highest concentration was found in the non-injection site muscle, at 0.945 mg kg^-1^ (*Rnext*). As *M* was greater than *Vmax*, Maximum Level of Exposure (*MLE*) for an individual vulture was calculated as *MLE* = *Rmax*\**Vmax* + *Rnext*\*(*M-Vmax*). Based on Galligan *et al*. (2022), we calculated the dose for individual vultures in mg kg^-1^ (*D1*) as *D1* = *MLE/W* and ml (*D2*) as *D2 = (MLE*W)/U*, where *U* was the concentration of nimesulide in the brand used (Nimovet; 100 mg ml^-1^). We calculated an MLE of 149.1 mg kg^-1^, a D1 of 18.64 mg kg^-1^ and a D2 of 1.50 ml for both vultures. Further details can be found in Galligan *et al*. (2022).

### 2.4 Blood sampling

We collected blood samples (4-5 ml) from each bird, by direct veno-puncture from the tarsal vein, at 0 hours (i.e., just before treatment), and at 2, 6, 12 and 24 hours after treatment, and serum and plasma separated. Serum tubes with clot activator and heparin-coated plasma tubes were used for collecting blood.

### 2.5 Extraction and measurement of nimesulide in vulture plasma

We extracted nimesulide from vulture plasma and tissue samples using a dilution method and measured concentrations of nimesulide in the acetonitrile using liquid chromatography tandem mass spectrometry (LC-MS/MS; Sciex API 6500+™) with electro spray ionization (ESI) and multiple reaction monitoring (MRM) in negative ionization mode.

Dimethyl sulfoxide (DMSO, 2.0 ml) was added to 2.12 mg of nimesulide in a 2.0 ml micro-centrifuge tube, mixed well and sonicated; 2 ml DMSO was added to 2.32 mg of nimesulide (D-5), which was used as an internal standard stock. Analyte working calibration standards (0.5 to 500 ng ml^-1^) and quality control samples (QC) of nimesulide were prepared in DMSO. An internal working standard solution was prepared by diluting 0.15 ml of the internal standard stock solution to 50 ml with acetonitrile to provide a concentration of 3.0 μg ml^-1^. This solution was mixed well and stored at 2 to 8°C. Calibration standards and quality control samples of nimesulide were prepared by spiking 47.5 μl of blank vulture plasma with 2.5 μl of the analyte working solution. Study samples (20 μl) were transferred to Eppendorf tubes and 10 μl of the internal working standard solution added. Samples were quenched with 200 μl of acetonitrile, vortexed and centrifuged at 14,000 rpm for 5 minutes at 4°C. 150 μl of supernatant was transferred to 1 ml vials for analysis by LC-MS/MS.

Further details of the methods for the extraction and measurement of nimesulide can be found in Appendix A of the Supplementary Information.

### 2.6. Measurement of serum constituents

We analysed serum samples to estimate the concentration of the following biochemical analytes, changes in concentrations of which are often a symptom of kidney failure: uric acid, creatinine, urea, total protein, albumin, alanine aminotransferase (ALT), aspartate aminotransferase (AST), alkaline phosphatase (ALP), sodium and potassium. Concentrations of serum biochemical analytes were measured using commercially available measurement kits (Coral Clinical Systems, Tulip Diagnostics) using a GENESYS 10UV spectrophotometer (Thermo Scientific).

### 2.7. The effect of nimesulide treatment on the concentrations of serum metabolites

We extracted metabolites of nimesulide from the serum of treated and control-group Himalayan Griffons, which were analysed using liquid chromatography mass spectrometry (LC-MS/MS). Further details can be found in the Supplementary Information, Appendix C.

### 2.8. Post-mortem examination of vultures

Post-mortem examination was carried out on any vulture that died during the safety testing experiment. We performed a necropsy, which focussed principally on the viscera because visceral gout is a typical finding in *Gyps* vultures killed by NSAID poisoning (Oaks *et al*., 2004; Swan *et al*., 2006a, Naidoo *et al*., 2010, Zorilla *et al*. 2014, Cuthbert *et al*. 2016, Nambirajan *et al*. 2021). Tissue samples were collected during the necropsy, fixed in 10% buffered formalin, embedded in paraffin wax and cut at 4 μm. After mounting, the sections were stained with Haematoxylin and Eosin (H&E), and silver nitrate / hydroquinone (De Galantha staining) to confirm the presence of urate crystals in the visceral organs.

### 2.9. Animal ethics

Permission to carry out experiments on vultures was granted by the Government of Haryana Forest Department, and the RSPB’s Animal Ethics Committee (EAC2016-02).

## 3. Results

### 3.1. Concentration of nimesulide in tissues

Pharmacokinetic profiles (Fig. 1) differed slightly between the two birds treated with nimesulide. Whereas nimesulide was rapidly absorbed up until 6 hours after dosing in bird X36, concentrations of the drug in the plasma of bird X33 fell between 2 and 6 hours. Subsequently, elimination of nimesulide occurred at a similar rate in both birds up until their deaths after 24 hours. The rate of elimination was slower than the rate of absorption. Nimesulide concentration in the plasma of the two birds assigned to the control group was zero.

**Figure 1.**
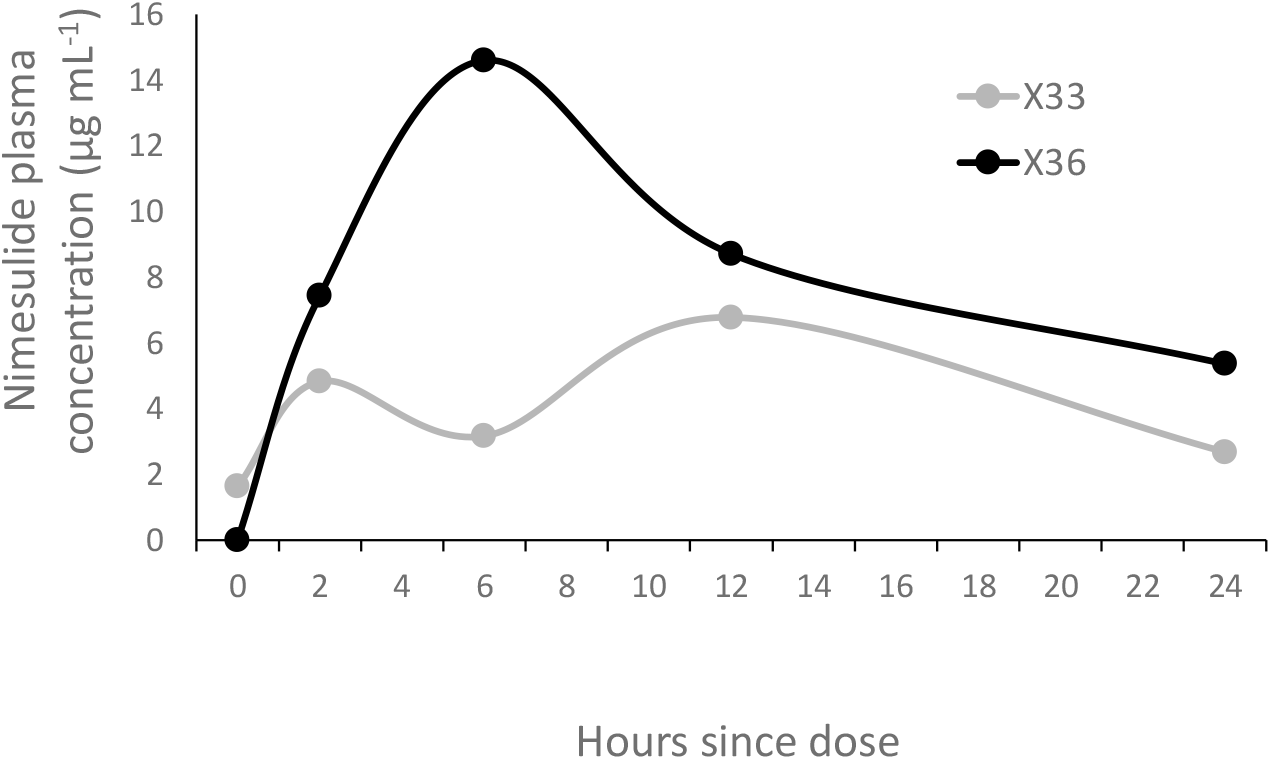
Pharmacokinetic profiles for the two Himalayan Griffons *Gyps himalayensis* treated with nimesulide.

### 3.2. Analysing samples for clinical pathology

Both treated vultures showed increases in serum uric acid concentrations (Fig. 2). Uric acid levels in the treated birds increased steadily until the birds died after 24 hours post-treatment. The concentrations just prior to death were 43-75-fold greater than the pre-treatment concentrations. Uric acid levels in control birds remained low throughout the experiment (Fig. 2).

**Figure 2.**
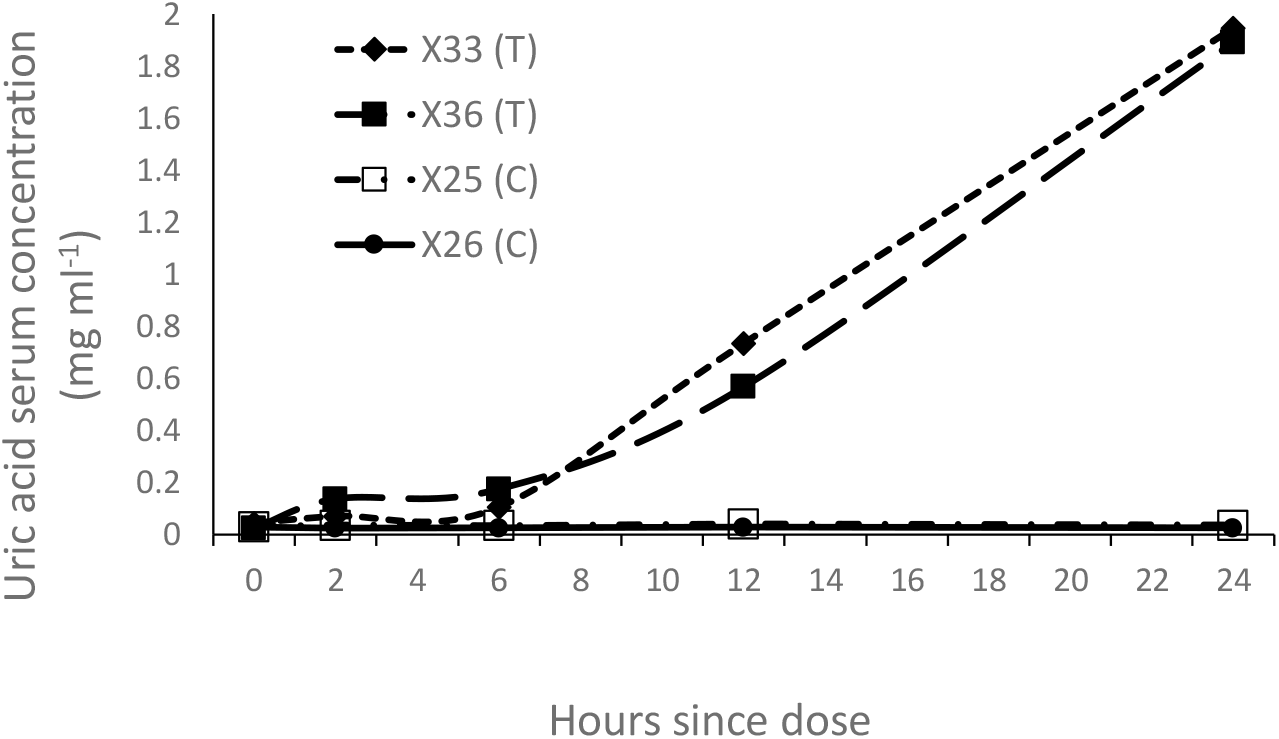
Change over time in plasma uric acid concentrations in two vultures treated with nimesulide (T), in comparison to two vultures assigned to a control group (C) and given a solution of benzyl alcohol / ethanol (the carrier solution for nimesulide in the Nimovet formulation).

There were large increases in serum potassium concentrations in treated birds (Fig. 3a). At 12 hours, potassium concentrations in treated vultures were 3-5 times higher than in controls, although levels stabilised between 12 and 24 hours; at 24 hours, the difference was only 0-2 times higher in treated birds. One bird (X33) showed a four-fold increase in aspartate aminotransferase (AST) up to 12 hours after treatment, with a slight decrease thereafter (Fig 3b). Although X36 also had relatively high AST levels, pre-treatment baseline levels for this bird were also the highest of all four birds. Creatinine levels were similar in treated and control birds up until 12 hours post-treatment, after which they increased in treated birds, remaining roughly stable in the controls (Fig. 3c). There were no other consistent differences between groups in changes in concentrations of the other analytes (Fig. S1).

**Figure 3.**
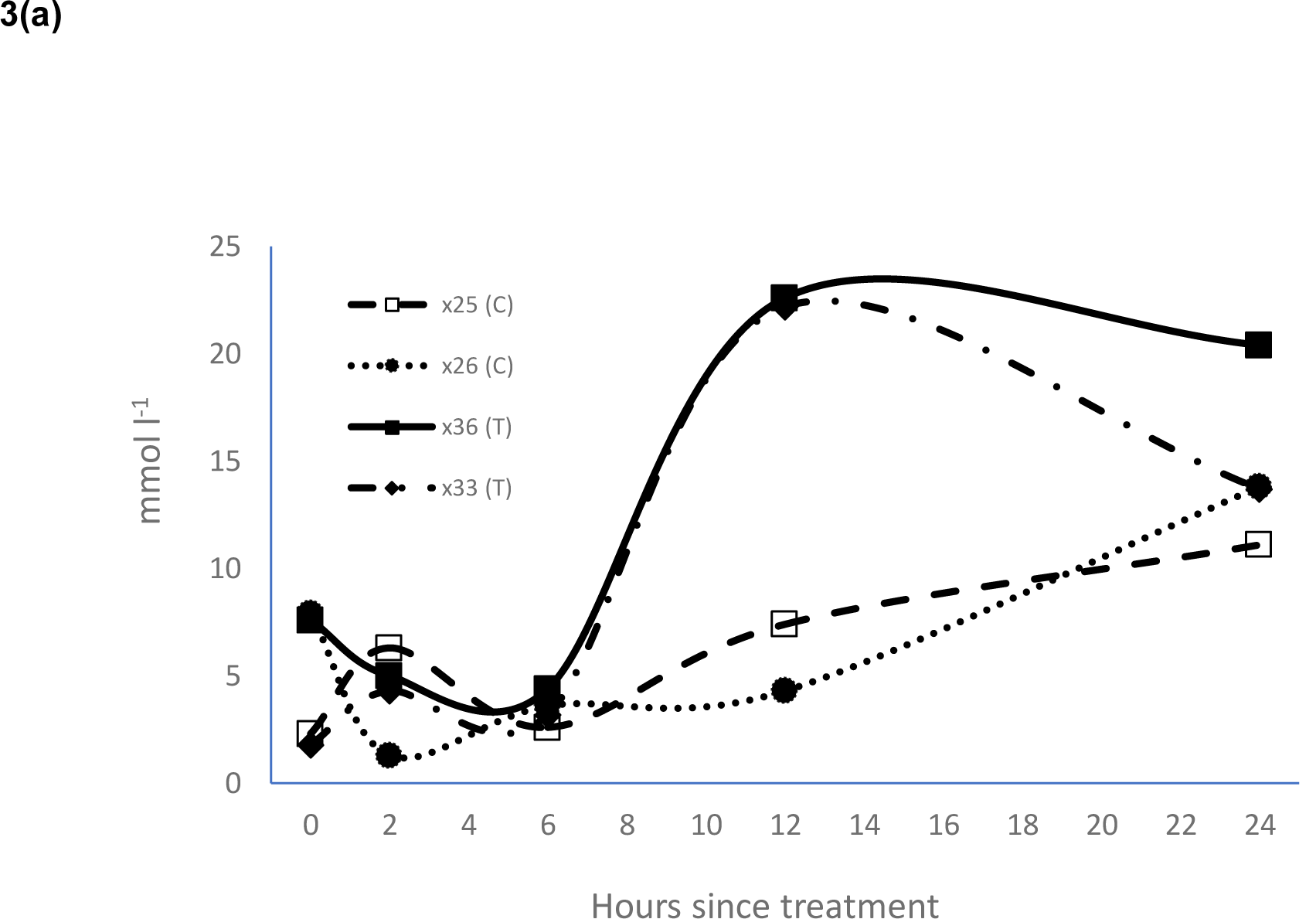
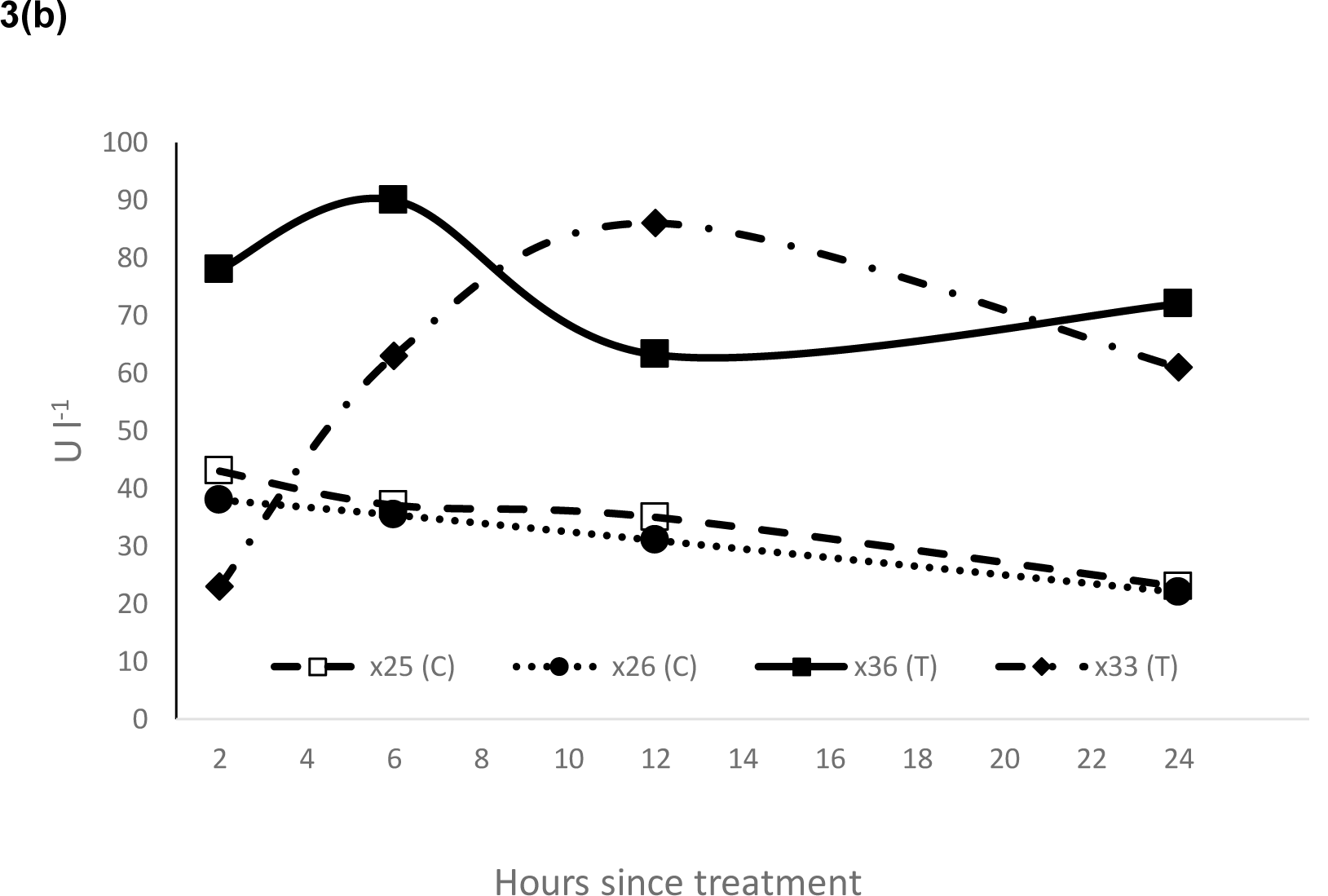
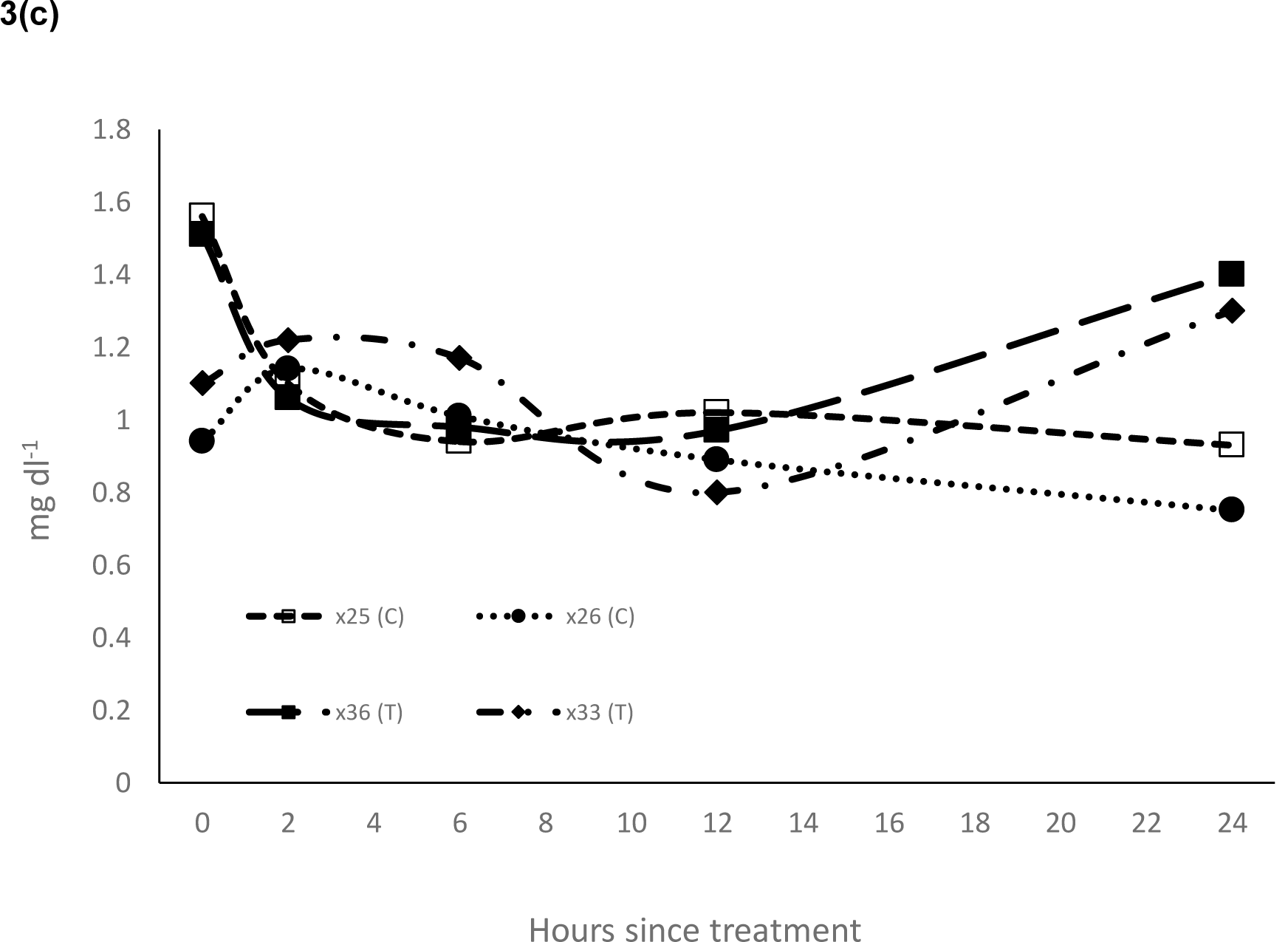
Change over time in plasma potassium (a), aspartate aminotransferase (AST) (b) and creatinine (c) concentrations in two vultures (T) treated with nimesulide in comparison to two vultures assigned to the control group (C) and given a solution of benzyl alcohol / ethanol (the carrier solution for nimesulide in the Nimovet formulation).

### 3.3. Changes in the concentration of serum metabolites

A total of 158 compounds were extracted from the serum of treated and control-group *G. himalayensis*. Of these, concentrations of 47 metabolites significantly increased after treatment with nimesulide compared to the control group, 33 significantly declined, and for the remaining 78 metabolites there was no change (Supplementary Information, Appendix C, Tables S2, S3 and S4).

### 3.4. Post-mortem results for treated G. himalayensis

Post-mortem analysis was carried out on the two *G. himalayensis* that died following oral gavage treatment with nimesulide. Systematic necropsy examination of both birds revealed deposition of chalky white urate crystals on the visceral organs (severe visceral gout; Fig. 4, a), caused by kidney failure. Staining confirmed the presence of urate crystals (tophi) in the visceral organs, i.e., kidney and liver (Fig. 4, b-d).

**Figure 4.**
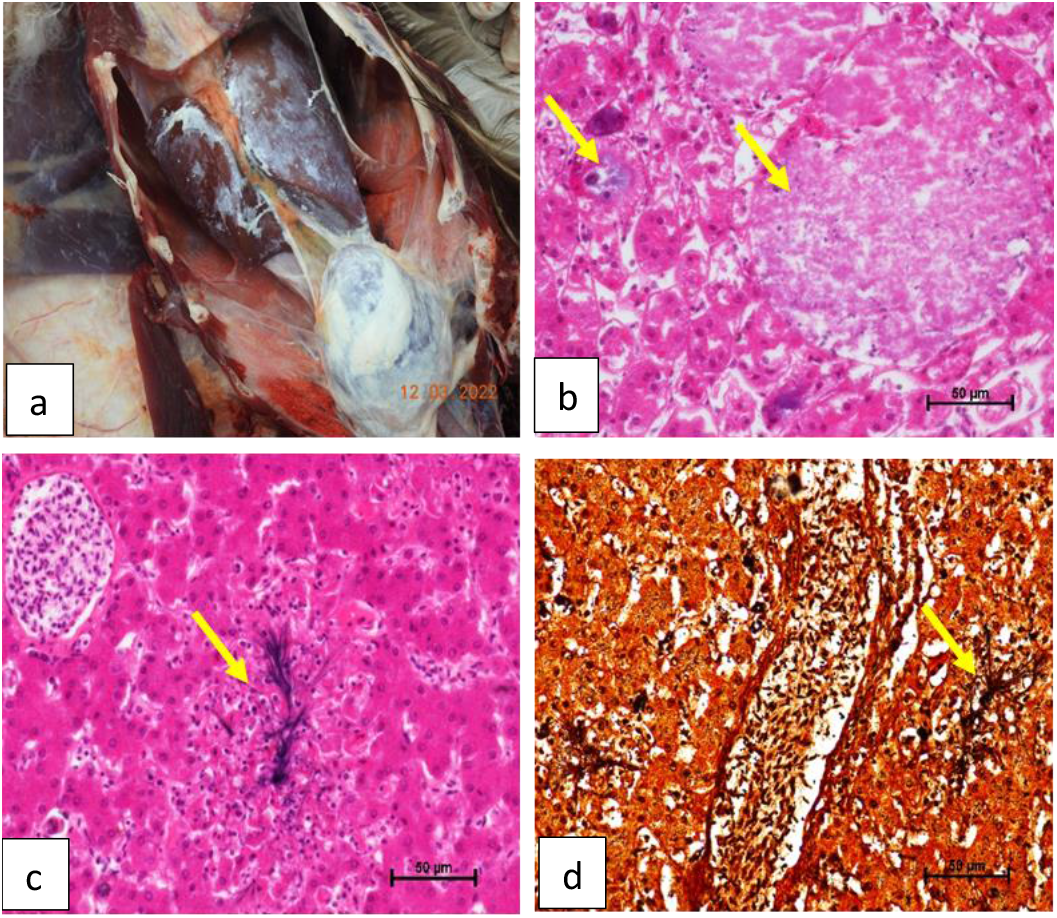
(a) Post-mortem image showing deposition of chalky white urate crystals on visceral organs. Histopathological images of (b) liver and (c) kidney tissues showing deposition of urate crystals, highlighted by Haematoxylin and Eosin (H&E) staining, and (d) silver nitrate / hydroquinone (De Galantha) staining (40 x magnification).

## 4. Discussion

The results of our study confirm that nimesulide is toxic to *Gyps* vultures. Both birds treated with the drug died around 24 hours after treatment, had elevated plasma levels of uric acid and showed post-mortem signs of severe visceral gout. These post-mortem outcomes have previously been recognised as indicating NSAID nephrotoxicity (Oaks *et al*., 2004; Swan *et al*., 2006a, Naidoo *et al*., 2010, Zorilla *et al*. 2014, Cuthbert *et al*. 2016, Nambirajan *et al*. 2021). In contrast, the two birds in the control group, which were given a dose of benzyl alcohol (the carrier solution of nimesulide), survived and showed no adverse symptoms. Several dead wild *Gyps bengalensis* in India have previously been found to have residues of nimesulide co-occurring with visceral gout and other symptoms of kidney failure in two independent studies (Cuthbert *et al*. 2016, Nambirajan *et al*. 2021). The toxicity of nimesulide to *Gyps* vultures was also established by safety testing experiments carried out on an African species, *G. coprotheres* (Galligan *et al*. 2022). Our study shows that nimesulide caused the deaths of two captive *G. himalayensis*. As far as we know, this is the only such experimental test of the toxicity of nimesulide conducted on an Asian species of *Gyps* vulture. These findings further highlight the need for veterinary use of nimesulide to be banned in India and other countries in Asia that are home to critically endangered species of vultures.

That nimesulide should be found to be as toxic to Himalayan Griffons as it was to Cape Griffons is not surprising given the sensitivity to NSAID toxicity shown by all species of *Gyps* vultures that have been tested. Although some unrelated species of scavenging birds appear to be tolerant to the effects of diclofenac, e.g., Turkey Vulture *Cathartes aura* (Rattner *et al*. 2008) and Pied Crow *Corvus albus* (Naidoo *et al*. 2011), individuals of all five *Gyps* species on which diclofenac was experimentally tested, i.e., *G. bengalensis* (Oaks *et al*. 2004), Eurasian Griffon *G. fulvus* and African White-backed Vulture *G. africanus* (Swan *et al*. 2006a), *G. coprotheres* (Naidoo *et al*. 2009) and *G. himalayensis* (Das *et al*. 2011)), all died showing the same typical symptoms of kidney failure, elevated uric acid levels and visceral gout. Additionally, carcasses of a sixth species, Indian Vulture, *Gyps indicus*, were regularly found throughout South Asia with the same symptoms (Shultz *et al*. 2004). It is thus likely that all species of *Gyps* vultures have the same sensitivity to NSAID toxicity (Swan *et al*. 2006b).

However, there is still a need for a better understanding of what other species are sensitive to NSAID toxicity. In south Asia, populations of two other species of vulture – Red-headed Vulture *Sarcogyps calvus* and Egyptian Vulture *Neophron percnopterus* – underwent declines of similar magnitude, and over a similar time period (although with a slight time-lag), to those of species of *Gyps* vulture (Cuthbert *et al*. 2006). They have also shown signs of recovery since the ban on veterinary use of diclofenac (Galligan *et al*. 2014). Therefore, it is quite possible that both these species are also sensitive to diclofenac toxicity, although no individuals of either species have been found with symptoms of NSAID toxicity, nor has experimental safety testing been carried out on these species. Other studies have, however, widened the range of scavenging species that are affected by NSAIDs, including Steppe Eagle *Aquila nipalensis* in India (Sharma *et al*. 2014). Furthermore, a nestling Cinereous Vulture *Aegypius monachus* in Spain recently became the first-known avian victim of diclofenac poisoning outside of Asia (Herrer-Villar *et al*. 2021). Also, experimental work has been carried out to investigate the toxicity of both diclofenac and nimesulide in Black Kites *Milvus migrans* (Farooq & Khan 2023a, b), common scavengers in Asia. Whereas being given diclofenac at relatively high doses resulted in the deaths of four out of six kites (Farooq & Khan 2023b), no birds died after being given nimesulide, although they did show the typical signs of nephrotoxicity (Farooq & Khan 2023a).

Changes in the concentrations of many metabolites in vultures treated with nimesulide (Supplementary Information, Appendix C), suggests inhibition of metabolic pathways and excretory processes in vultures. Although these results are preliminary due to the small sample size, in future, with larger sample sizes, we may be able to use metabolomics to identify the biomarkers of NSAID toxicity. Also, by comparing metabolic differences between vulture-toxic and vulture-safe NSAIDs, such methods could be used as a non-lethal alternative to current safety testing experiments.

Nimesulide is increasingly being sold in pharmacies for the treatment of cattle in both India and Nepal (Galligan *et al*. 2021). Although in India figures vary by state, up to 37.3% of pharmacies sampled within the State of Haryana in 2017 offered the drug; while in Nepal, up to 13.7% of pharmacies in the western Terai in 2016 did so (Galligan et al., 2021). Given the evidence of the toxicity of nimesulide to vultures from several studies, there is an urgent need for the drug to be banned in India. This will not only benefit vultures in India, but across the wider South Asian region, too.

## CRediT authorship contribution statement

**Karikalan Mathesh:** Formal analysis, Investigation, Data Curation, Writing – Review & Editing. **Kesavan Manickam:** Formal analysis, Investigation. **John Mallord:** Writing – Original Draft, Project administration. **K. Mahendran:** Formal analysis, Investigation. **Asok Kumar M:** Formal analysis, Investigation. **Krishna Chutia:** Formal analysis, Investigation. **Debasish Saikia:** Formal analysis, Investigation. **Beena V:** Formal analysis, Investigation. **Sree Lakshmi P:** Formal analysis, Investigation. **Nikita Prakash:** Resources, Project administration. **Rohan Shringarpure:** Formal analysis, Investigation. **Abhijit Pawde:** Formal analysis, Investigation. **Rhys Green:** Conceptualization, Methodology, Writing – Review & Editing. **Vibhu Prakash:** Conceptualization, Methodology, Writing – Review & Editing.

## Declaration of competing interest

The authors declare that they have no known competing financial interests or personal relationships that could have appeared to influence the work reported in this paper.

## Acknowledgments

We are grateful to BNHS and the Director, ICAR-IVRI for their general support, and to Eurofins Advinus Ltd, Bengalaru, India for extraction and quantification of nimesulide in plasma samples. The work was funded by the Ministry of Environment, Forest and Climate Change of the Government of India, Haryana State Forest Department, and the RSPB. The funding bodies had no involvement in the execution of the experiment nor the production of this manuscript.

## Notes

### Competing Interest Statement

The authors have declared no competing interest.

