## Appendix A for "Experimental safety testing confirms that the NSAID nimesulide is toxic to *Gyps* vultures in India"

**SUPPLEMENTARY INFORMATION**

**Appendix A: Methodology to estimate the concentration of nimesulide in vulture plasma samples**

| Name of compound | Analyte | Internal standards |
| --- | --- | --- |
|  | Nimesulide | Nimesulide-D5 |
| Molecular weight of free compound (base / acid) | 308.31 | 313.34 |
| Diluent | DMSO | |
| Calibration curve range / Internal standard Working Concentration | 0.5 to 500 ng ml^-1^ | 3.0 µg ml^-1^ |
| Chromatographic conditions: | | |
| Mobile phase | Pump A: 0.1 % formic Acid in 5 mM ammonium acetate solution  Pump B: Acetonitrile | |
| Gradient conditions | Binary Gradient: Time (min) \| % B concentration; 0.01 \| Start; 0.50 \| 25; 1.50 \| 85; 3.50 \| 85; 3.60 \| 25; 5.00 \| Stop | |
| Column (make) | Zorbax XDB C8, 50*4.6 mm, 5µ | |
| Injection volume (µL) | 10 µL | |
| Flow rate (mL/min) | 1 | |
| Run time (min) | 5 | |
| Sample cooler temperature (°C) | 5 | |
| Column oven temperature (°C) | 40 | |
| Rinsing solution | 50% Acetonitrile | |
| Plasma Sample preparation: Protein precipitation method | | |
| CC & QC preparation: An aliquot (47.5 µL) of blank plasma was spiked with 2.5 µL of analyte working solution.  Sample Treatment Procedure: 20 µL of CC / QC / Study samples were aliquoted into pre-labeled eppendorff tubes and 10 µL of internal working standard solution was added. Samples were quenched with 200 µL of Acetonitrile and vortexed. All the samples were centrifuged at 14000 rpm for 5 minutes at 4^0^C. 150 µL of supernatant was transferred into inserts kept in 1 mL vials and capped with polyethylene plugs and analyzed in LC-MS/MS. | | |

Mass spectrometric condition :

| Instrument ID | API 6500+ LC-MS/MS | | |
| --- | --- | --- | --- |
| Mass parameters | Analyte:  Nimesulide | IS:  Nimesulide-D5 | |
| MRM transitions | 307.2 → 228.8  307.2 → 79.2 | 312.1 → 234.1 | |
| Resolution – Q1 | Unit | Unit | |
| Resolution – Q3 | Unit | Unit | |
| Declustering potential (DP) Volts | -51 | -45 | |
| Entrance potential (EP) | 10 | | |
| Collision energy (CE) Volts | -21  -44 | | -25 |
| Collision cell exit potential  (CXP) Volts | -12 | | -12 |
| Ionisation / Polarity | Negative | | |
| Dwell time (milliseconds) | 500 | | |
| Ionisation Source | ESI | | |
| IS | -4500 | | |
| Collision gas (CAD) | Medium | | |
| Curtain gas (CUR) | 35 | | |
| GS1 | 65 | | |
| GS2 | 70 | | |
| Temperature | 500 °C | | |
