## Appendix B, Table S1 for "Experimental safety testing confirms that the NSAID nimesulide is toxic to *Gyps* vultures in India"

**Concentrations of constituents of blood serum of Himalayan Griffon vultures in relation to time since dosing with nimesulide**

**Table S1.** Change in plasma concentrations of various biochemical analytes in two vultures treated with nimesulide (T) in comparison to two vultures assigned to a control group (C) and given a solution of benzyl alcohol / ethanol (the carrier solution for nimesulide in the Nimovet formulation).

|  | **Uric acid (mg dl^-1^)** | | | | | **Creatinine (U l^-1^)** | | | | | **Total Protein (g dl^-1^)** | | | | |
| --- | --- | --- | --- | --- | --- | --- | --- | --- | --- | --- | --- | --- | --- | --- | --- |
| **Bird** | **0h** | **2h** | **6h** | **12h** | **24h** | **0h** | **2h** | **6h** | **12h** | **24h** | **0h** | **2h** | **6h** | **12h** | **24h** |
| **X25 (C)** | 3.07 | 3.7 | 3.5 | 4.12 | 3.6 | 1.56 | 1.1 | 0.94 | 1.02 | 0.93 | 2.2 | 2.3 | 3.9 | 2.1 | 2.7 |
| **X26 (C)** | 2.9 | 2.5 | 2.6 | 2.81 | 2.6 | 0.94 | 1.14 | 1.01 | 0.89 | 0.75 | 4.3 | 2 | 3.7 | 1.8 | 2.4 |
| **X36 (T)** | 2.5 | 13.7 | 17.5 | 56.8 | 189.1 | 1.51 | 1.06 | 0.98 | 0.97 | 1.4 | 1.8 | 1.9 | 3.5 | 2.2 | 2.3 |
| **X33 (T)** | 4.5 | 7.1 | 10.6 | 73.5 | 194.6 | 1.1 | 1.22 | 1.17 | 0.8 | 1.3 | 1.6 | 1.7 | 3.3 | 1.9 | 2.3 |
|  | **Albumin (g dl^-1^)** | | | | | **AST (U l^-1^)** | | | | | **ALT (U l^-1^)** | | | | |
| **Bird** | **0h** | **2h** | **6h** | **12h** | **24h** | **0h** | **2h** | **6h** | **12h** | **24h** | **0h** | **2h** | **6h** | **12h** | **24h** |
| **X25 (C)** | 1 | 1.1 | 1.13 | 1.16 | 1.4 | 56 | 43 | 37.1 | 35 | 23 | 75 | 75.1 | 69.1 | 72.7 | 46.5 |
| **X26 (C)** | 0.87 | 0.94 | 0.99 | 0.66 | 1.5 | 32.4 | 38 | 35.4 | 31 | 22 | 73 | 74.6 | 69 | 71.4 | 46.1 |
| **X36 (T)** | 0.96 | 0.91 | 0.95 | 0.68 | 1.3 | 83 | 78 | 90 | 63.2 | 72 | 73 | 73 | 69.7 | 70 | 42.8 |
| **X33 (T)** | 1.1 | 0.8 | 0.87 | 0.59 | 1.42 | 21 | 23 | 63 | 86 | 61 | 73.8 | 72.1 | 69.1 | 69.4 | 42.8 |
|  | **Sodium (mmol l^-1^)** | | | | | **Potassium (mmol l^-1^)** | | | | | **ALP (U l^-1^)** | | | | |
| **Bird** | **0h** | **2h** | **6h** | **12h** | **24h** | **0h** | **2h** | **6h** | **12h** | **24h** | **0h** | **2h** | **6h** | **12h** | **24h** |
| **X25 (C)** | 87 | 82 | 107 | 68 | 80.1 | 2.3 | 6.3 | 2.6 | 7.4 | 11.1 | 2.4 | 2.4 | 2.4 | 3 | 3 |
| **X26 (C)** | 77 | 61.3 | 88 | 52 | 66 | 7.9 | 1.3 | 3.6 | 4.3 | 13.8 | 2.5 | 2.4 | 3.1 | 2.9 | 3.1 |
| **X36 (T)** | 73 | 43 | 46 | 56.2 | 68 | 7.6 | 5 | 4.4 | 22.6 | 20.4 | 2.4 | 2.5 | 2.9 | 3.1 | 3.2 |
| **X33 (T)** | 99.1 | 62 | 61.3 | 63 | 64 | 1.8 | 4.3 | 3.2 | 22.2 | 13.7 | 2.42 | 2.45 | 2.8 | 3.2 | 3.3 |
