## Appendix C, Tables S2-S4 for "Experimental safety testing confirms that the NSAID nimesulide is toxic to *Gyps* vultures in India"

**SUPPLEMENTARY INFORMATION**

**Appendix C**

**The effect of nimesulide treatment on the concentrations of serum metabolites**

*Methods*

**Sample Collection and Preparation**

Plasma samples were stored at − 80 °C until analysis. Before analysis samples were thawed at 4 °C, and homogenized in an ice bath with a Vortex oscillator for 60 s. 200 μl 80% methanol was added to 30 μl of homogenized sample and mixed for 60 s. They were centrifuged at 10,000 × g for 5 min at 4 °C to remove precipitates and the supernatants were collected for analysis. 100 ul supernatant was filtered with 0.22u PTFE filter and transferred into 150 μl glass insert in a 1.5 ml amber glass vial and analyzed by UHPLC-MS/MS.

**Liquid chromatography mass spectrometry analysis (LC/MS)**

The LC/MS experiments were carried out using the Dionex UltiMate 3000 UHPLC (Thermo Scientific, MA) system equipped with a binary pump and an auto sampler, coupled with Thermo Q-EXACTIVE PLUS Mass spectrometer operating in positive (ESI +) and negative (ESI−) electrospray ionization mode (Thermo Fisher Scientific, Sunnyvale, CA, USA), separately. The chromatographic separation was performed with a Thermo Hypersil Gold C18 column (100 mm × 2.1 mm, 1.9 μm particle size). Separation was achieved with solvent A (Water + 0.1% (v/v) formic acid) and solvent B (acetonitrile + 0.1% (v/v) formic acid) with the following gradient at a flow rate of 0.3 ml/min: 0 min (5% B), 0–8 min (5–95% B), 8–11 min (95% B), 11–12 min (95 – 5 % B) and 12–15 min (5% B). The injection volume was 15 μl and the analysis time was 15 min.

The mass spectrometer parameters in this experiment were as follows:

1. Voltage: 4 kV
2. Capillary temperature: 350 °C
3. Resolution (First Full Scan):1,40,000
4. Scan range: 100–1500 m/z; AGC target:5e5.
5. The resolution of the secondary data dependency scan (Full MS / dd-MS2) was 35000 and scan range of 200-2000 m/z; AGC target:5e4.

***Quality Control***

For validating the stability of the analysis process, quality control (QC) sample was first prepared by mixing from each sample (10 μl). QC was injected once (In both Positive and Negative mode) before, middle and at the end of the batch process to monitor the stability of sample preparation and instrument.

There was also a blank run in-between the samples to clean the analytical column and tubings.

***Data Processing***

All samples were processed, and RAW files generated were analyzed with Compound Discoverer (CD) - 3.2. For compound consolidation, Mass Tolerance was set at 5 ppm and Retention Time (RT) was to 0.2 min. Metabolite annotation was performed through ChemSpider and mzCloud databases. ChemSpider database search was performed with BioCyc, HMDB, ChEBI, MMDB, MOLPORT, Nature Chemical Biology, PubMed, Royal Society of Chemistry, Serum Metabolome database and KEGG database.

The results of analysis were exported with following filtration criteria: delta PPM range from -5 ppm to 5, Background is False, Name is fully annotated and for which all annotations have MS/MS.

Both positive and negative mode analysis was performed separately, which was combined at later stage and duplicates were removed. Statistical analysis was performed on redundant list of metabolites. Intensity values were log2 transformed, z-score normalised, and T-test was applied on comparisons reported in the Results.

*Results*

A total of 158 compounds were extracted from the serum of treated and control-group Himalayan Griffons. Of these, concentrations of 47 metabolites significantly increased after treatment with nimesulide compared to the control group, 33 significantly declined, and for the 78 metabolites there were no changes (Tables S2, S3 and S4).

*Discussion*

Changes in the concentrations of many metabolites in vultures treated with nimesulide (Supplementary Information, Appendix C), suggests inhibition of metabolic pathways and excretory processes in vultures. Although these results are preliminary due to the small sample size, in the future, with larger sample sizes, we may be able to use metabolomics to identify the biomarkers of NSAID toxicity. Also, by comparing metabolic differences between vulture-toxic and vulture-safe NSAIDs, LC-MS-based metabolomics could be used as a non-lethal alternative to current safety testing experiments.

Table S2. List of serum metabolites whose concentrations significantly increased after birds were given a dose of nimesulide.

| **Metabolite** | **Formula** | **logFC** | ***P*** |
| --- | --- | --- | --- |
| (+)-neocroalbidine | C18 H29 N O6 | 1.88 | 0.003 |
| (6E,8E)-3-Hydroxy-10-methoxy-4,9-dimethyl-10-oxo-6,8-decadienoic acid | C13 H20 O5 | 3.61 | 0.001 |
| 1,5-DAN | C10 H10 N2 | 2.82 | 0.004 |
| 1,5-Di-O-acetyl-2,3,4,6-tetra-O-methylhexitol | C14 H26 O8 | 1.29 | 0.023 |
| 2-({1-[4,6-Bis(dimethylamino)-1,3,5-triazin-2-yl]-1H-1,2,4-triazol-3-yl}sulfanyl)acetamide | C11 H17 N9 O S | 2.39 | 0.003 |
| 2-(1-Acetyl-3-piperidinyl)acetamide | C9 H16 N2 O2 | 1.98 | 0.008 |
| 2-Ethyl-2-({2-hydroxy-3-[4-methoxy-2-(2-methyl-2-propanyl)phenoxy]propyl}amino)-1,3-propanediol | C19 H33 N O5 | 6 | 0.013 |
| 2-Methyl-2-propanyl [(2S)-1-hydroxy-3-(3,4,5-trimethoxyphenyl)-2-propanyl]carbamate | C17 H27 N O6 | 3.35 | 0.001 |
| 2-Methyl-2-propanyl 2-{[(2-methoxyethyl)sulfamoyl]methyl}-4-morpholinecarboxylate | C13 H26 N2 O6 S | 2.60 | 0.004 |
| 2-Methyl-2-propanyl 3-{1-[3-(methylsulfonyl)propyl]-2-piperidinyl}propanoate | C16 H31 N O4 S | 3.33 | 0.004 |
| 2-Methyl-2-propanyl methyl(2-{methyl[(pentylsulfonyl)acetyl]amino}ethyl)carbamate | C16 H32 N2 O5 S | 3.48 | <0.0001 |
| 2,2'-(propane-1,3-diylbis(oxy))bis-1,3,2-dioxaborinane | C9 H18 B2 O6 | 1.01 | 0.028 |
| 3-[(1R,3S)-1-Hydroxy-3-(2-hydroxyethoxy)-7-azaspiro[3.5]non-7-yl]-N,N-dimethyl-1-propanesulfonamide | C15 H30 N2 O5 S | 3.94 | <0.0001 |
| 4-(Methylsulfonyl)-N-[3-(1H-tetrazol-5-yl)propyl]-1,4-diazepane-1-carboxamide | C11 H21 N7 O3 S | 1.82 | 0.001 |
| 4-Methyl-N-[3-methyl-1-oxo-1-(1,4,7,10-tetraoxa-13-azacyclopentadecan-13-yl)-2-butanyl]benzenesulfonamide | C22 H36 N2 O7 S | 4.49 | <0.0001 |
| 4-Morpholinyl[4-(4-morpholinylsulfonyl)-1-piperazinyl]methanone | C13 H24 N4 O5 S | 3.05 | 0.0003 |
| 4,4'-({6-[(2-Aminoethyl)amino]-1,3,5-triazine-2,4-diyl}diimino)di(1-butanol) | C13 H27 N7 O2 | 1.73 | 0.005 |
| 5-[(2,6-Dibromo-4-chlorophenyl)hydrazono]-2,4,6(1H,3H,5H)-pyrimidinetrione | C10 H5 Br2 Cl N4 O3 | 1.85 | 0.014 |
| 5-Nitroxystearic acid | C18 H35 N O5 | 1.73 | 0.005 |
| 8-Amino-7-oxononanoic acid | C9 H17 N O3 | 0.87 | 0.029 |
| Bis(2-methyl-2-propanyl) (2R,3R)-2-hydroxy-3-[(2-morpholinylcarbonothioyl)oxy]succinate | C17 H29 N O7 S | 3.42 | 0.001 |
| bis(tetrazolyl)amine | C2 H3 N9 | 1.49 | 0.011 |
| Boc-10-Adc-OH | C15 H29 N O4 | 1.43 | 0.014 |
| BZ-LYS-OH | C13 H18 N2 O3 | 3.44 | 0.0001 |
| Diethyl 2,3-pyridinedicarboxylate | C11 H13 N O4 | 2.63 | 0.001 |
| Echimidine | C20 H31 N O7 | 4.59 | 0.001 |
| Ethyl [(1,1-dioxidotetrahydro-3-thiophenyl)(ethyl)amino](oxo)acetate | C10 H17 N O5 S | 2.56 | 0.001 |
| Ethyl 5-cyclohexyl-4,6-dioxo-1,3a,4,5,6,6a-hexahydropyrrolo[3,4-c]pyrazole-3-carboxylate | C14 H19 N3 O4 | 2.17 | 0.003 |
| ethynyl cyclopentadienone | C7 H4 O | 3.23 | 0.002 |
| GV9100000 | C8 H12 O | 2.73 | 0.003 |
| Leucylproline | C11 H20 N2 O3 | 0.87 | 0.029 |
| Methyl [2-(2,3-dihydro-1H-indol-1-yl)-4-oxo-1,4,5,6-tetrahydro-5-pyrimidinyl]acetate | C15 H17 N3 O3 | 2.82 | 0.002 |
| Methyl N-[(4,5-dimethoxy-2-methylphenyl)sulfonyl]-2-methylnorvalinate | C16 H25 N O6 S | 4.98 | <0.0001 |
| N-(3-Methoxyphenyl)-N'-(3-methylbutyl)ethanediamide | C14 H20 N2 O3 | 3.56 | 0.003 |
| N-(6-Oxaspiro[4.5]dec-9-yl)methanesulfonamide | C10 H19 N O3 S | 4.34 | 0.001 |
| N-[(2-Methoxyadamantan-2-yl)methyl]methanesulfonamide | C13 H23 N O3 S | 2.22 | 0.021 |
| NI3910000 | C4 H6 N4 O | 2.38 | 0.006 |
| Nitrosoheptamethyleneimine | C7 H14 N2 O | 3.88 | 0.001 |
| NP-001608 | C13 H16 O5 | 2.95 | 0.005 |
| NP-012017 | C13 H22 O5 | 2.09 | 0.014 |
| Pentoxifylline | C13 H18 N4 O3 | 8.30 | <0.0001 |
| Phomalone | C13 H18 O5 | 1.80 | 0.011 |
| Pirbuterol | C12 H20 N2 O3 | 1.24 | 0.014 |
| PLK | C17 H32 N4 O4 | 1.65 | 0.004 |
| Protopine | C20 H19 N O5 | 1.14 | 0.008 |
| tert-Butyl [2-(2-aminoethoxy)ethyl]carbamate | C9 H20 N2 O3 | 1.18 | 0.018 |
| tert-Butyl 4-[2-(hydroxyamino)-2-iminoethyl]piperazine-1-carboxylate | C11 H22 N4 O3 | 2.58 | 0.003 |

Table S3. List of serum metabolites whose concentrations significantly decreased after birds were given a dose of nimesulide.

| **Name of Metabolite** | **Formula** | **logFC** | **P.Value** |
| --- | --- | --- | --- |
| (15Z,19R)-25-Amino-22-hydroxy-22-oxido-17,21,23-trioxa-22lambda~5~-phosphapentacos-15-en-19-yl docosanoate | C43 H86 N O7 P | -2.80 | 0.0003 |
| (3R,6R)-N-(2-Acetamidoethyl)-6-[(4-{[benzyl(methyl)amino]methyl}-1H-1,2,3-triazol-1-yl)methyl]quinuclidine-3-carboxamide | C24 H35 N7 O2 | -1.47 | 0.005 |
| (Nitroimino)dimethanol | C2 H6 N2 O4 | -2.70 | 0.0008 |
| (Z)-N-[2-(Hexadecylsulfanyl)ethyl]-1-(3-nitrophenyl)methanimine | C25 H42 N2 O2 S | -1.31 | 0.025 |
| 1-deoxymethylsphinganine | C17 H37 N O | -1.06 | 0.029 |
| 2-Bromo-1,3,4-oxadiazole | C2 H Br N2 O | -2.06 | 0.011 |
| 2-Trifluoromethylpyrrolidine | C5 H8 F3 N | -1.24 | 0.021 |
| 2-vinylquinazolin-4-ol | C10 H8 N2 O | -3.10 | 0.001 |
| 2,2'-[(2R,5S)-1,4-Dioxane-2,5-diylbis(methyleneimino)]diethanol | C10 H22 N2 O4 | -2.57 | 0.024 |
| 2,3,4-Tri-O-methyl-L-threo-pentitol | C8 H18 O5 | -4.18 | 0.011 |
| 2,6-Difluorostyrene | C8 H6 F2 | -2.62 | 0.001 |
| 3-(Docosanoyloxy)-2-[(9Z,12Z)-9,12-nonadecadienoyloxy]propyl (4Z,7Z,10Z,13Z,16Z)-4,7,10,13,16-docosapentaenoate | C66 H114 O6 | -1.14 | 0.049 |
| 3-Fluor-2-methylpyridin | C6 H6 F N | -2.04 | 0.01 |
| 4-Amino-N'-[(Z)-(4-amino-1,2,5-oxadiazol-3-yl)(hydroxyimino)methyl]-1,2,5-oxadiazole-3-carbohydrazonamide | C6 H8 N10 O3 | -2.25 | 0.038 |
| 4-Imino-1,2,3-butanetriol | C4 H9 N O3 | -2.20 | 0.014 |
| 5-Aminovaleric acid | C5 H11 N O2 | -1.86 | 0.003 |
| 6-azido-2(S)-hydroxyhexanoic acid | C6 H11 N3 O3 | -2.266 | 0.029 |
| 781825 | C10 H8 N2 O2 | -1.776 | 0.006 |
| Allylsulfanyl-acetic acid | C5 H8 O2 S | -2.016 | 0.020 |
| Bilirubin | C33 H36 N4 O6 | -5.416 | <0.0001 |
| Bis(4-chlorophenyl) hydrogen phosphate | C12 H9 Cl2 O4 P | -1.296 | 0.017 |
| Elacytarabine | C27 H45 N3 O6 | -1.20 | 0.013 |
| Fmoc-Hcit-OH | C22 H25 N3 O5 | -1.56 | 0.033 |
| Hexadecyl 2,5,7-trinitro-9-oxo-9H-fluorene-4-carboxylate | C30 H37 N3 O9 | -4.64 | 0.007 |
| L-(-)-Methionine | C5 H11 N O2 S | -2.01 | 0.020 |
| L-Methionine sulfoxide | C5 H11 N O3 S | -2.10 | 0.003 |
| MFCD00144495 | C2 H3 D2 I | -1.03 | 0.018 |
| N,N-dimethylsulfamide | C2 H8 N2 O2 S | -2.53 | 0.007 |
| Nicotinamide | C6 H6 N2 O | -1.81 | 0.046 |
| PAF C-16 Carboxylic Acid | C26 H52 N O9 P | -1.53 | 0.042 |
| PAPA NONOate | C6 H16 N4 O2 | -2.22 | 0.004 |
| Pentachloroethane | C2 H Cl5 | -1.76 | 0.004 |
| Tetrafluoro(selenoxo)molybdenum | F4 Mo Se | -1.86 | 0.004 |

Table S4. List of serum metabolites whose concentrations did not significantly change after birds were given a dose of nimesulide.

| **Name of Metabolite** | **Formula** | **logFC** | **P.Value** |
| --- | --- | --- | --- |
| (11Z)-eicoseneoylcarnitine | C27 H51 N O4 | -0.99 | 0.195 |
| (1E)-1-(2-Furyl)-1,4-pentadien-3-one | C9 H8 O2 | -0.58 | 0.175 |
| (1R,4S,5R,6S)-5,6-Dihydroxy-2-azabicyclo[2.2.1]heptan-3-one | C6 H9 N O3 | -1.65 | 0.221 |
| (2R)-2-({(2R,3R,4S,5R,6R)-4-{[(2R,3R,4R,5S,6R)-3-Acetamido-6-({[({(2R,3R,4S,5R,6R)-4-{[(2R,3R,4R,5S,6R)-3-acetamido-4,5-dihydroxy-6-(hydroxymethyl)tetrahydro-2H-pyran-2-yl]oxy}-6-[(1R)-1-carboxy-2-hydroxyethoxy]-3,5-dihydroxytetrahydro-2H-pyran-2-yl}methoxy)(hydroxy)phosphoryl]oxy}methyl)-4,5-dihydroxytetrahydro-2H-pyran-2-yl]oxy}-6-[({[(5-aminopentyl)oxy](hydroxy)phosphoryl}oxy)methyl]-3,5-dihydroxytetrahydro-2H-pyran-2-yl}oxy)-3-hydroxypr" | C39 H69 N3 O33 P2 | -0.14 | 0.849 |
| (2S,3R)-2-[(13Z)-13-Docosenoylamino]-3-hydroxyoctadecyl 5-acetamido-3,5-dideoxy-6-[(1R,2R)-1,2,3-trihydroxypropyl]-beta-L-threo-hex-2-ulopyranonosyl-(2->3)-[5-acetamido-3,5-dideoxy-6-[(1R,2R)-1,2,3-trihydroxypropyl]-beta-L-threo-hex-2-ulopyranonosyl-(2->3)-beta-D-galactopyranosyl-(1->3)-2-deoxy-2-(2-oxopropyl)-beta-D-galactopyranosyl-(1->4)]-beta-D-galactopyranosyl-(1->4)-beta-D-glucopyranoside" | C89 H157 N3 O39 | -1.04 | 0.057 |
| (2Z,2'Z)-2,2'-(1,4-Cyclooctanediylidene)dihydrazinecarboxamide | C10 H18 N6 O2 | -0.54 | 0.692 |
| (4S)-4-[(2-Hydroxyhexadecanoyl)oxy]-4-(trimethylammonio)butanoate | C23 H45 N O5 | -1.64 | 0.107 |
| (5-hydroxy-3-methyl-1-phenyl-1H-pyrazol-4-yl)(5-methyl-2-phenyl-2H-1,2,3-triazol-4-yl)methanone | C20 H17 N5 O2 | 1.56 | 0.278 |
| (9xi,11beta,14xi,17alpha)-11,17-Dihydroxy-3-oxoandrosta-1,4-diene-17-carboxylic acid | C20 H26 O5 | 0.06 | 0.931 |
| (Dimethylamino)[(2,5-dioxo-1-pyrrolidinyl)oxy]-N,N-dimethylmethaniminium | C9 H16 N3 O3 | -0.77 | 0.385 |
| 1-Boc-4-piperidinemethanol | C11 H21 N O3 | -1.36 | 0.120 |
| 1-Methylhistidine | C7 H11 N3 O2 | 0.17 | 0.678 |
| 1,1,1,3-Tetrabromoacetone | C3 H2 Br4 O | -1.75 | 0.065 |
| 1,3-Dimethoxy-2-propanyl methanesulfonate | C6 H14 O5 S | -2.13 | 0.175 |
| 1,5-Naphthalenediamine | C10 H10 N2 | -0.78 | 0.351 |
| 10-[(4-Nitrophenyl)methyl]-1,4,7,10-tetraazacyclododecane-1,4,7-triacetamide | C21 H34 N8 O5 | 1.00 | 0.201 |
| 11-Aminoundecanoic acid | C11 H23 N O2 | -2.10 | 0.246 |
| 1760709 | C10 H24 N2 O3 | -0.76 | 0.244 |
| 1H-1-Benzosilole | C8 H8 Si | -0.34 | 0.406 |
| 2-(4-Methoxy-2,2,6,6-tetramethyl-1-piperidinyl)ethyl 4-oxopentanoate | C17 H31 N O4 | 0.03 | 0.943 |
| 2-{[4-(4-Methylphenyl)-5-phenyl-4H-1,2,4-triazol-3-yl]sulfanyl}-N-(2,4,6-tribromophenyl)acetamide | C23 H17 Br3 N4 O S | -0.58 | 0.148 |
| 2-Amino-9-[3-azido-4-hydroxy-3-(hydroxymethyl)butyl]-3,9-dihydro-6H-purin-6-one | C10 H14 N8 O3 | -1.01 | 0.527 |
| 2-Bromoacrylamide | C3 H4 Br N O | -0.52 | 0.646 |
| 2-Butyltetrahydrofuran | C8 H16 O | -1.02 | 0.335 |
| 2-Methyl-2-propanyl (3aS,6S,6aS)-2,2-dimethyl-6-tetradecyltetrahydrofuro[3,4-d][1,3]oxazole-3(2H)-carboxylate | C26 H49 N O4 | -0.94 | 0.169 |
| 2,4,8-Triamino-3-oxooctanoic acid | C8 H17 N3 O3 | 0.34 | 0.323 |
| 3-(Dimethylamino)-3-[1-(2-methoxyethyl)-1H-tetrazol-5-yl]-N-(3-methylbutyl)-1-piperidinecarboxamide | C17 H33 N7 O2 | -0.60 | 0.347 |
| 3-(Pentachlorophenoxy)propanoic acid | C9 H5 Cl5 O3 | -0.86 | 0.201 |
| 3-{2-[2-(2,5-Dioxo-2,5-dihydro-1H-pyrrol-1-yl)ethoxy]ethoxy}-N,N-bis{2-[2-(2-propyn-1-yloxy)ethoxy]ethyl}propanamide | C25 H36 N2 O9 | -0.65 | 0.187 |
| 3-Mesityl-1-(2-methylcyclohexyl)-1-[(1-propyl-1,2,3,4-tetrahydro-6-quinolinyl)methyl]thiourea | C30 H43 N3 S | -1.40 | 0.085 |
| 3,3-Dimethyl-1,5-dioxaspiro(5.5)undecan-9-one | C11 H18 O3 | -2.00 | 0.183 |
| 3,6-Dibromo-1,5-dihydro-4H-pyrazolo[3,4-d]pyrimidin-4-one | C5 H2 Br2 N4 O | -2.37 | 0.068 |
| 3b-Hydroxy-5-cholenoic acid | C24 H38 O3 | -0.36 | 0.470 |
| 4-{[(2-Methoxyethyl)(octanoyl)amino]methyl}phenyl methanesulfonate | C19 H31 N O5 S | 0.61 | 0.543 |
| 4-Hydroxytetrahydro-2H-thiopyran-4-carboxylic acid | C6 H10 O3 S | -0.61 | 0.206 |
| 5-chlorotryptamine | C10 H11 Cl N2 | -0.61 | 0.206 |
| 5,6-Octadiene-1,3-diyne | C8 H6 | -0.83 | 0.130 |
| 6-Amino-5-(2,2-diethoxyethyl)-2-mercapto-4-pyrimidinol | C10 H17 N3 O3 S | -0.65 | 0.400 |
| 6-Methylthiochroman-4-one | C10 H10 O S | -0.41 | 0.678 |
| 8-[(tert-Butoxycarbonyl)amino]-1,4-dioxaspiro[4.5]decane-8-carboxylic acid | C14 H23 N O6 | -0.11 | 0.870 |
| 9-Octadecenamide | C18 H35 N O | -0.65 | 0.195 |
| a-(Aminomethyl)-4-morpholineethanol | C7 H16 N2 O2 | -0.07 | 0.902 |
| Acetyl-L-carnitine | C9 H17 N O4 | -0.75 | 0.120 |
| Acetyl-methylcholine | C8 H17 N O2 | -0.48 | 0.184 |
| D-(-)-Glutamine | C5 H10 N2 O3 | -0.15 | 0.804 |
| Decanoylcarnitine | C17 H33 N O4 | -0.31 | 0.499 |
| Dichlorogermane | H2 Cl2 Ge | -1.17 | 0.184 |
| Dihydrothymine | C5 H8 N2 O2 | 0.17 | 0.678 |
| DL-Glutamine | C5 H10 N2 O3 | 1.10 | 0.152 |
| Docosahexaenoic acid ethyl ester | C24 H36 O2 | -0.36 | 0.470 |
| Ethyl N~2~-acetyl-L-argininate | C10 H20 N4 O3 | 0.34 | 0.323 |
| FMOC-CHA-OH | C24 H27 N O4 | -1.52 | 0.183 |
| GLPG0187 | C29 H37 N7 O5 S | -1.74 | 0.158 |
| hypaphorine | C14 H18 N2 O2 | -1.24 | 0.120 |
| Indole | C8 H7 N | -0.05 | 0.953 |
| iodonitrotetrazolium | C H I N5 O2 | -0.57 | 0.186 |
| L-Histidine | C6 H9 N3 O2 | -0.78 | 0.309 |
| L-Leucyl-L-leucyl-L-phenylalanylglycyl-L-tyrosyl-L-alanyl-L-valyl-L-tyrosyl-L-valine | C54 H77 N9 O12 | -2.07 | 0.181 |
| L-Phenylalanine | C9 H11 N O2 | -0.58 | 0.175 |
| L-Pyroglutamic acid | C5 H7 N O3 | -0.44 | 0.499 |
| Methyl (3,4,5-triethoxy-2-nitrophenyl)acetate | C15 H21 N O7 | -0.80 | 0.206 |
| Methyl 4-O-benzyl-6-deoxy-alpha-L-mannopyranoside | C14 H20 O5 | 0.80 | 0.201 |
| MFCD00227369 | C18 H19 Cl3 | 0.55 | 0.274 |
| MFCD09841385 | C14 H9 D9 N4 O3 | -0.96 | 0.193 |
| ML-10 | C9 H15 F O4 | -0.64 | 0.241 |
| N-[(4-Hexylcyclohexyl)carbonyl]methionine | C18 H33 N O3 S | 1.18 | 0.146 |
| N-[2-(Diethylamino)ethyl]-5-nitro-4,6-pyrimidinediamine | C10 H18 N6 O2 | -2.89 | 0.152 |
| N-[3-(Dimethylamino)propyl]-5-[(3R)-1,2-dithiolan-3-yl]pentanamide | C13 H26 N2 O S2 | -1.41 | 0.232 |
| N-[4-(Benzyloxy)phenyl]-4-(3-methylphenoxy)butanamide | C24 H25 N O3 | -1.46 | 0.181 |
| N,N'-1,2-Ethanediylbis[2-(2,4-dimethoxyphenyl)-4-quinolinecarboxamide] | C38 H34 N4 O6 | -1.79 | 0.190 |
| N,N-Bis(cyanomethyl)-2-(3,4-dimethoxyphenyl)acetamide | C14 H15 N3 O3 | 1.30 | 0.120 |
| N~5~-(Diaminomethylene)-N-[(2R)-6-(dimethylamino)-2-hexanyl]-L-ornithinamide | C14 H32 N6 O | -1.08 | 0.252 |
| N6-Methoxy-9H-purine-2,6-diamine | C6 H8 N6 O | -0.90 | 0.400 |
| N6,N6,N6-Trimethyl-L-lysine | C9 H20 N2 O2 | 0.37 | 0.25 |
| Oleonitrile | C18 H33 N | -0.65 | 0.195 |
| Palmitoylcarnitine | C23 H45 N O4 | 0.09 | 0.875 |
| Stearoyl glutamic acid | C23 H43 N O5 | -0.78 | 0.242 |
| UV9702000 | C8 H12 N4 | -0.31 | 0.730 |
